# Quantification of individual dataset contributions to prediction accuracy in cooperative learning

**DOI:** 10.1101/2025.04.16.649215

**Authors:** David Dechantsreiter, Rachel S. Kelly, Christoph Lange, Jessica Lasky-Su, Georg Hahn

## Abstract

We consider cooperative learning, a recently proposed technique to leverage the predictive power of several datasets (also called data views) to improve the prediction accuracy of an outcome of interest. Cooperative learning uses a Lasso-type penalty to fit several datasets to a given outcome, while a so-called agreement penalty enforces that the predictions made by the individual datasets agree. We are interested in the question of whether the predictive power of each individual dataset can be quantified using cooperative learning. In this work, we demonstrate that certain trace plots, analogously to the ones for the classic Lasso, allow one to quantify the predictive power of each individual dataset. Importantly, this allows one to detect datasets which do not carry any predictive power on the outcome. In an experimental study, we quantify the predictive power of three real datasets in the context of the Childhood Asthma Management Program (CAMP), with the three datasets containing information on epidemiological variables, metabolites, and clinical data, respectively.

## 1 Introduction

One important aspect in the age of precision medicine pertains to estimating the susceptibility of an individual to a given disease. To this end, given input data are fitted to some outcome of interest. For instance, polygenic risk scores [1] and integrated risk models [2] quantify the aggregated genetic risk for a disease or trait, and have been shown to identify people at high risk for certain diseases [3, 4, 5].

Oftentimes, several datasets might be predictive of an outcome of interest. However, predicting an outcome separately with each dataset might result in deviating or even contradictory forecasts. In these cases, cooperative learning allows one to combine the predictions of several data sources in a meaningful and consistent way [6]. To be precise, cooperative learning takes as inputs *m* ∈ ℕ datasets which are thought to be predictive of some outcome, with the only requirement being that the datasets agree in the number of rows. The number of predictors in each dataset (columns) can be chosen freely. A Lasso-type penalty [7] enforces sparse predictions for each of the *m* datasets, while a so-called agreement penalty ensures that the predictions are consistent, in the sense that after fitting all datasets yield roughly the same prediction.

Since the input datasets are arbitrary, their contribution to the final prediction is unknown a priori. We are interested in the question whether the predictive power of each individual dataset can be quantified using cooperative learning. Since in the cooperative learning framework, the individual predictions are added up, we observe that comparing the final prediction made by cooperative learning to the ones made with each input dataset alone allows one to quantify their proportional contribution to the final prediction. This contribution can then be plotted as a function of the agreement penalty. We demonstrate that the resulting trace plot allows one to quantify the predictive power of each individual dataset. This trace plot is analogous to the one commonly computed for ridge regression or the Lasso, showing when certain predictors enter the model [8]. Importantly, the proposed cooperative trace plots allow one to detect datasets which do not carry any predictive power on the outcome.

We demonstrate the proposed methodology using an application of clinical relevance using datasets from the Childhood Asthma Management Program (CAMP) [9, 10]. We aim to predict the FEV1 (Forced Expiratory Volume in 1 second), a common spirometric measure of lung function. The three datasets used for prediction contain information on epidemiological variables (such as age and sex), metabolites, and clinical data, respectively. By computing cooperative trace plots for the prediction, we are able to show what proportion of the prediction stems from each dataset, and how this changes as the agreement penalty increases.

This article is structured as follows. Section 2 gives an overview of cooperative learning and introduces the proposed methodology. The experimental study is presented in Section 3. The article concludes with a discussion in Section 4.

Throughout the article, we denote with *X*^(*S*)^ the submatrix consisting of the rows of a matrix *X* that are indexed in some set *S* ⊆ {1, …, *n*}, where *n* is the number of rows of *X*. The compliment operation, that is the subset of rows of *X* which are not in *S* is denoted with *X*^(*−S*)^. Analogously, we denote with *y*^(*S*)^ the subset of entries of a vector *y* indexed in *S*, and with *y*^(*−S*)^ the subset of entries not in *S*.

## 2 Methods

In this section, we briefly review the concept of cooperative learning (Section 2.1) before introducing trace plots (Section 2.2).

### 2.1 Cooperative learning

We are given *m* ∈ ℕ datasets labeled *X*_1_, …, *X*_*m*_ having dimensions 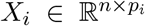, where *i* ∈ {1, …, *m*}. We aim to predict an outcome *y* ∈ ℝ^*n*^ of length *n* ∈ ℕ. Importantly, cooperative learning requires each dataset *X*_*i*_ to hold information about all *n* ∈ ℕ outcomes, but the number of predictors *p*_*i*_ in *X*_*i*_ can differ between the datasets.

Two straightforward ways to use the *m* datasets for the prediction of *y* are called *early fusion* and *late fusion*. In early fusion, we concatenate all datasets by columns into one matrix *X* = [*X*_1_, …, *X*_*m*_] having dimensions *n* × *p*, where 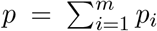. We then predict *y* from *X* using any common prediction framework, for instance linear regression, Lasso regression [7], or machine learning. Importantly, the disadvantage of early fusion consists in the fact that for big datasets, the fusing of all data might not be feasible. In late fusion, we carry out *m* separate predictions using each *X*_*i*_, where *i* ∈ {1, …, *m*}, resulting in *m* predictions *y*_1_, …, *y*_*m*_. Since there is no guarantee that those *m* predictions agree or are consistent, we are faced with the problem to fuse those into one final prediction. For instance, one common strategy would be to average all predictions. However, this approach has the disadvantage that (1) there is no obvious way to combine predictions, and (2) it is usually the case that after combining (for instance, averaging) the *m* predictions, we are left with a final answer that is different from each individual prediction and thus not supported by any of the *m* datasets.

To alleviate these issues, cooperative learning allows one to both carry out the estimation and to enforce consistency among the estimates. We suppose that the prediction of the outcome with each dataset *X*_*i*_ is performed using some function *f*_*i*_, which usually depends on some unknown parameters *β*_*i*_, where *i* ∈ {1, …, *m*}. Notably, the functions *f*_*i*_ can be chosen freely, meaning that each prediction can be computed with (potentially) a different technique (such as linear regression for one prediction, and a convolutional neural network for another). Cooperative learning (see eq. (20) in [6]) considers the objective function

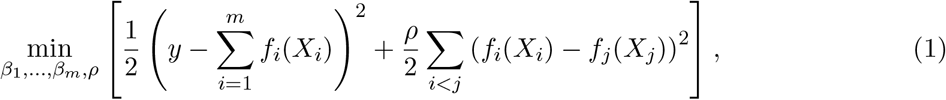

where *ρ >* 0 is the *agreement parameter* which controls the weight with which the predictions are enforced to agree. The selection of *ρ* can be carried out in various ways. If cross-validation is used, *ρ* is tuned externally and would not be fitted in eq. (1). The same applies if *ρ* is chosen a priori based on domain knowledge. Last, *ρ* can be fitted in a data adaptive way by including it in the optimization, which is the case in eq. (1). After the optimization of eq. (1) is completed, we obtain a fitted vector 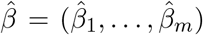 containing the fitted parameters 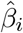 for each of the functions *f*_*i*_, where *i* ∈ {1, …, *m*}.

As noted in [6], the cooperative learning framework includes both early and late fusion as special cases. In particular, late fusion is obtained from eq. (1) as *ρ* → 0, since in this case no agreement on the prediction will be enforced. On the other hand, *ρ* → ∞ causes the predictions *f*_*i*_(*X*_*i*_) in eq. (1) to agree for all *i* ∈ {1, …, *m*}, however this might come at the expense of a reduced prediction quality.

### 2.2 Trace plots

The choice of the agreement penalty *ρ* in eq. (1) is not unique. Similarly to the Lasso penalty, commonly denoted as *λ >* 0 [7], the agreement penalty *ρ* can be fitted via cross-validation. However, analogously to the Lasso variable trace plots, which show when variables enter the Lasso model, it is natural to ask how the prediction made by cooperative learning varies with *ρ*. We aim to address this question by investigating how the prediction of each dataset *X*_*i*_ changes as a function of *ρ*, where *i* ∈ {1, …, *m*}.

To this end, we propose the following methodology. First, we split the data into a training proportion *α* ∈ (0, 1) and validation proportion 1−*α*. This is done by selecting a random subset *S* ⊆ {1, …, *n*} of size ⌊*αn*⌋. We then split all datasets *X*_1_, …, *X*_*m*_ into a training subset 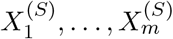 and a validation subset 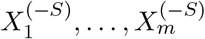. Similarly, the outcome vector *y* is split into a training subset *y*^(*S*)^ and a validation subset *y*^(*−S*)^.

Second, we use the training proportion of our data to fit a cooperative learning model of the type of eq. (1) for a fixed choice of *ρ*. This yields a fitted vector of parameters 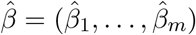 for each of the *m* functions *f*_1_, …, *f*_*m*_.

Third, using the unseen validation data, we evaluate the fit in two ways. On the one hand, we make a prediction with the full cooperative model, that is we make a prediction for *y*^(*−S*)^ given by

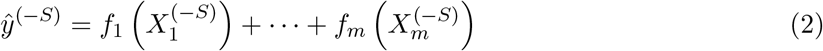

using the fitted coefficients 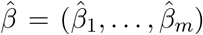 for *f*_1_, …, *f*_*m*_. Second we predict *y*^(*−S*)^ using each dataset individually, resulting in *m* predictions ranging from 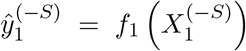 to 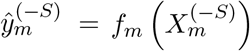. By comparing each of the 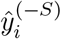 for *i* ∈ {1, …, *m*} with the full prediction 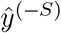 we aim to quantify how much each dataset contributed to the prediction.

Fourth, we propose to measure the contribution of each dataset with some sort of summary statistic. There is no unique choice to achieve this, as several metrics might be applicable depending on the type of data being predicted. If the outcome *y* ∈ ℝ^*n*^ is a vector, then each 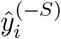 for *i* ∈ {1, …, *m*} can be compared with the full prediction 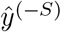 using vector norm, vector correlation, etc. Even simpler, we can compute the componentwise quotient 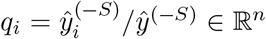 which will indicate what proportion of each entry in 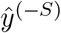 stems from the prediction with dataset 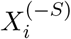. Importantly, this also allows one to gauge if some prediction underestimates or overestimates entries in 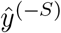, which are subsequently corrected with another prediction stemming from one of the other datasets. Finally, to visualize *q*_*i*_ ∈ ℝ^*n*^ for each prediction *i* ∈ {1, …, *m*}, we can plot all entries in *q*_*i*_ or some summary of it such as the mean or median.

The aforementioned computation can be repeated for several values of *ρ*. Plotting the quotients *q*_*i*_ for each dataset *i* ∈ {1, …, *m*} as a function of *ρ* will yield a cooperative trace plot, which we investigate further in the experimental results (Section 3).

## 3 Experimental results

This section presents our experimental results, starting with an overview of the experimental setting and the datasets we use (Section 3.1). We then demonstrate two examples of the trace plots in Section 3.2. In Section 3.3, we investigate the behavior of the trace plots in several scenarios in which some of the dataset carry predictive power of the outcome and some do not.

### 3.1 Setting and dataset

We employed data from the Childhood Asthma Management Program (CAMP). CAMP recruited children aged 5-12 years old with mild to moderate asthma. All children completed a protocol including questionnaires, blood collection, clinical testing and spirometry at recruitment and at regular in-person study visits across the five years of the trial. This current analysis uses data and blood samples collected at the end of the trial when the children had a median age of 13.1 years (range 9.1 to 17.2 years) [9].

Global plasma metabolomic profiling for CAMP was generated as part of the TOPMed initiative [10]. Four complementary ultrahigh-performance liquid-chromatography tandem mass spectrometry (LC-MS) platforms at the Broad Institute provided metabolomic profiles for CAMP. Three non-targeted and an additional targeted LC-MS methods using high resolution, accurate mass (HRAM) profiling to measure: (1) polar and nonpolar lipids; (2) free fatty acids, bile acids, and metabolites of intermediate polarity; and (3) polar metabolites including amino acids, acylcarnitines, and amines. An additional targeted LC-MS profiling method measured intermediary metabolites including purines and pyrimidines, and acyl CoAs.

We grouped the aforementioned data sources into three datasets containing information on *n* = 807 subjects. The first dataset *X*_1_ ∈ ℝ^807*×*4^ contains four epidemiological variables: age (range as above), sex (encoded as 1 for male and 2 for female), standing height in cm, and weight in kg. The second dataset *X*_2_ ∈ ℝ^807*×*92^ contains 92 measured metabolite levels (a precise list is given in the supplementary material). The third dataset *X*_3_ ∈ ℝ^807*×*5^ contains clinical data, specifically a collection of skin test results (number of core tests, number of positive core tests, number of skin tests, number of positive skin tests). The outcome for all *n* subjects is FEV1 measured as a positive real number. We randomly split the set of subjects using the proportions 70% − 30% to establish a training-validation split, yielding a training set (565 subjects) and a validation set (242 subjects).

### 3.2 An illustration of the trace plot

We apply cooperative learning to the three datasets of Section 3.1 while varying the agreement penalty *ρ* from 0 to 0.01 in steps of 0.001. For each choice of *ρ*, we fit the cooperative learning model of eq. (1) to the training datasets, thus obtaining three fitted coefficient vectors 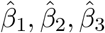 (see Section 2.1). We then use the fitted coefficients to evaluate both the full prediction 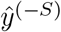 (where *S* denotes the set of indices belonging to the subjets in the training set) and the individual predictions 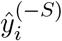 for each dataset *i* ∈ {1, 2, 3} to calculate the quotients *q*_*i*_ ∈ ℝ^2^42. The quotients *q*_*i*_ ∈ ℝ^2^42 quantify the proportion of prediction attributed to each dataset *i* ∈ {1, 2, 3}.

Figure 1 (left) shows the entries of quotient vector *q*_*i*_ for each dataset *i* ∈ {1, 2, 3} as a function of *ρ*. When summarizing each quotient vector *q*_*i*_ with its median, we obtain the smoothed version of the same plot depicted in Figure 1 (right). Three observations are noteworthy. First, the entries cluster in (remarkably) condensed point clouds for each dataset and each value of *ρ*. Second, as *ρ* increases, the proportion of the contribution of each dataset to the final prediction converges to 1*/*3. This is as expected, since *ρ* → ∞ in eq. (1) will force all predictions to agree, and 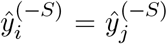 for each *i* ≠ *j* necessarily implies that each vector contributes 1*/*3 to the final prediction vector. However, as will be shown, this is only the case here since all three datasets are selected in such a way as to carry predictive power of the outcome. Third, the predictive proportion of each dataset can vary substantially for small values of *ρ*.

**Figure 1.**
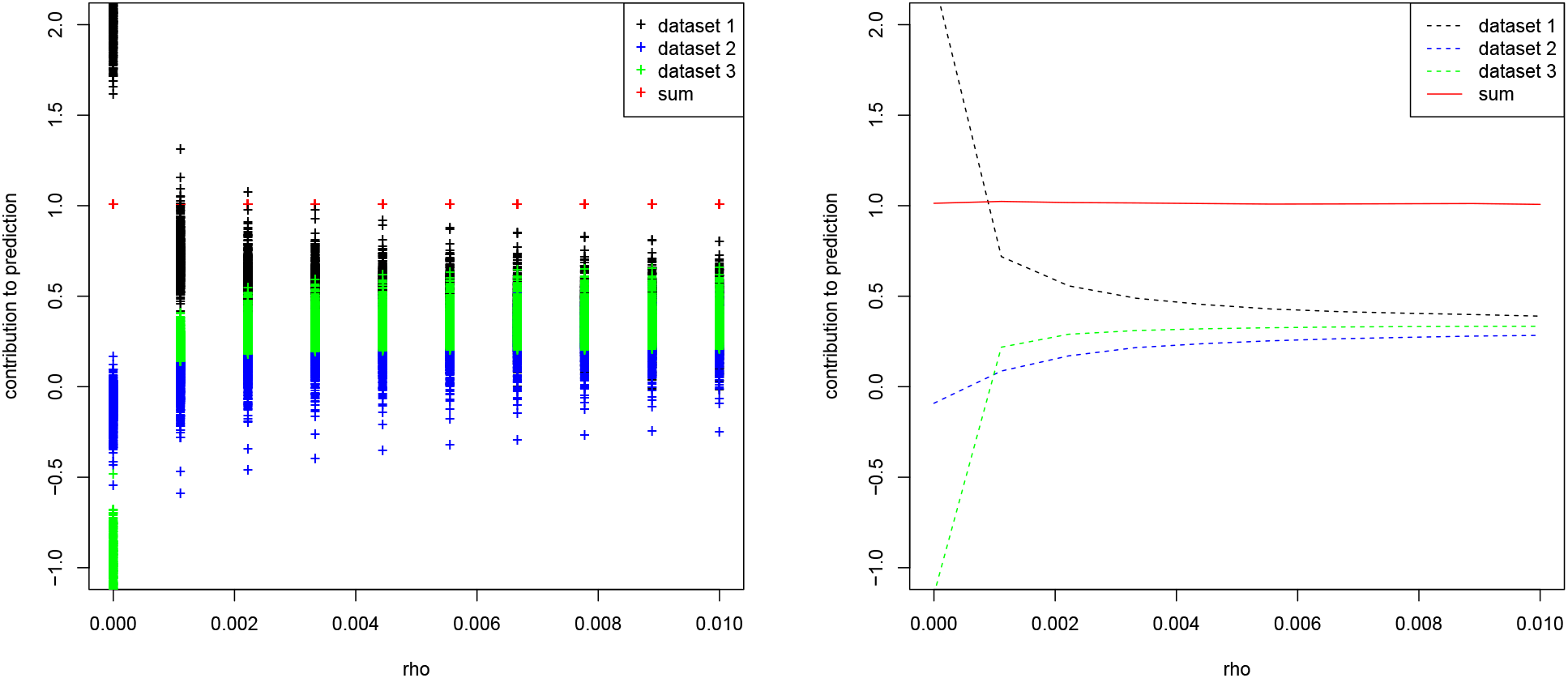
FEV1 dataset. Entries of the quotient vector *q*_*i*_ for each dataset *i* ∈ {1, 2, 3} (left) and median of each quotient vector *q*_*i*_ (right) as a function of the agreement penalty *ρ*.

While Figure 1 showed the proportions 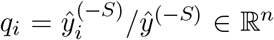, we might also be interested in looking at the predictive contribution of each dataset *i* ∈ {1, 2, 3} compared with the truth, that is 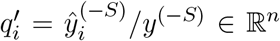, where *y*^(*−S*)^ denotes the withheld true outcome for the validation set. This is shown in Figure 2. We observe that the plot is visibly unchanged from Figure 1 (right), indicating that the prediction 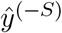 with cooperative learning is very close to the withheld truth *y*^(*−S*)^.

**Figure 2.**
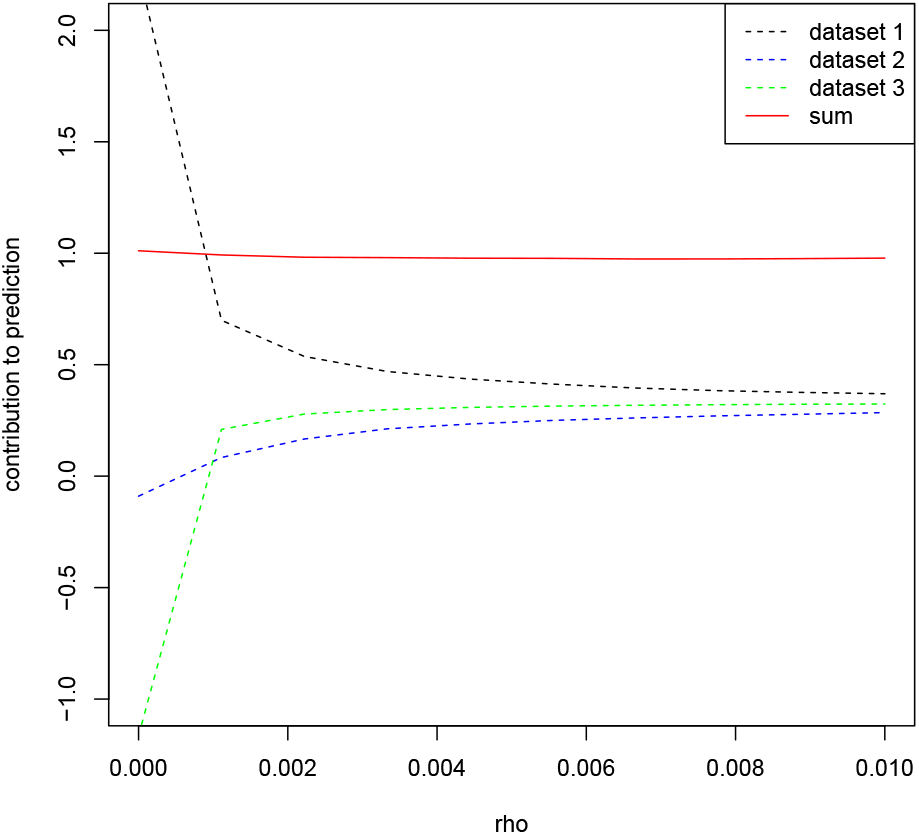
FEV1 dataset. Median of the entries of 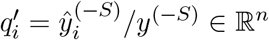, where *y*^(*−S*)^ denotes the withheld true outcome for the validation set.

### 3.3 Behavior for datasets carrying no predictive power

The three datasets selected in Section 3.1 all carry some predictive power of the outcome. In this section, we repeat the computation of the trace plots, but replace one or more of the three datasets with Gaussian noise. To be precise, we generate three random matrices *Y*_1_ ∈ ℝ^807*×*4^, *Y*_2_ ∈ ℝ^807*×*92^, and *Y*_3_ ∈ ℝ^807*×*5^ having the same dimensions as *X*_1_, *X*_2_, *X*_3_. Each matrix *Y*_*i*_ contains entries drawn from a Normal distribution *N* (0, 1) with mean zero and standard deviation one. Before fitting the cooperative learning model, we replace one or more dataset *X*_*i*_ with *Y*_*i*_, where *i* ∈ {1, 2, 3}.

Figure 3 shows the result of this experiment. Interestingly, we observe that if a dataset contains no predictive power, its contribution in cooperative learning to the full prediction remains at zero irrespective of the agreement penalty *ρ*. In turn, the two other datasets with predictive power converge to an equal contribution of 1*/*2 as *ρ* increases. This observation holds true when replacing any of the three datasets with Gaussian noise. Even when replacing two datasets with noise, the fitting process in cooperative learning assign negligible coefficients, resulting in the one remaining dataset contributing all entries to the final prediction, which is expected. In short, the cooperative trace plots allow one to quantify the predictive power of each dataset.

**Figure 3.**
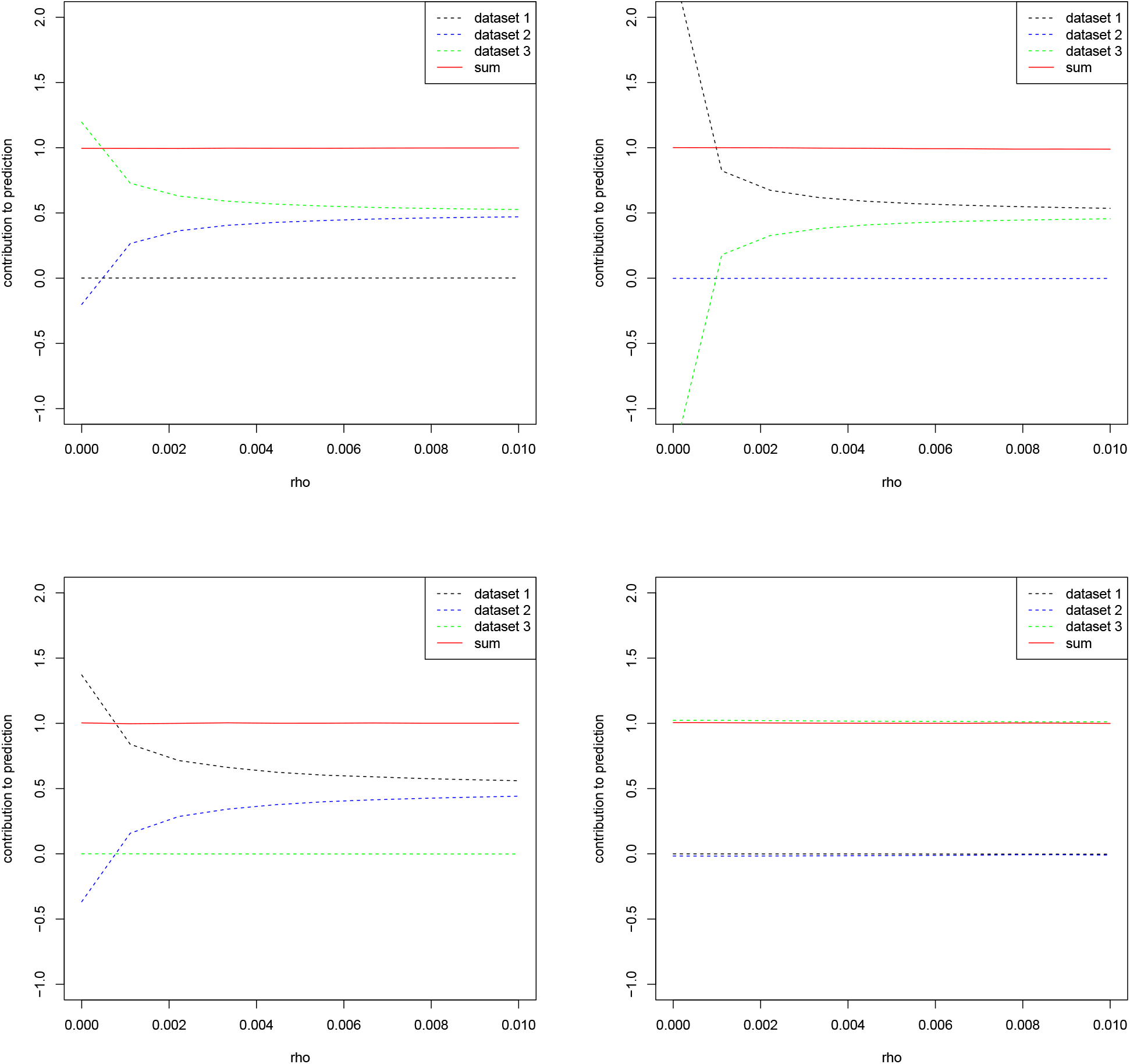
FEV1 dataset. Median of the quotient vector *q*_*i*_ for the three datasets of Section 3.1 when replacing either the first dataset with Gaussian noise (top left), the second dataset (top right), the third dataset (bottom left), or the first and second datasets (bottom right).

## 4 Discussion

In this contribution, we propose to compute trace plots for cooperative learning, a recently proposed statistical technique to leverage the predictive power of several datasets on some outcome of interest. The trace plots are computed by summarizing the componentwise contribution of each dataset’s individual prediction to the combined prediction made by cooperative learning. Our key findings can be summarized as follows.

First, the componentwise (entrywise) contribution of each dataset’s individual prediction to the full prediction is consistent across the subjects in the prediction set. Second, when summarizing the componentwise contributions with their median, we observe that the contribution of each dataset can vary substantially for small values of the agreement penalty *ρ*, and that the metrics converge towards an equal contribution as *ρ* increases, as expected. Third, for datasets with no predictive power, the cooperative trace plots show a contribution of zero, while the datasets with predictive power converge towards an equal contribution as *ρ* increases.

In summary, the cooperative trace plots allow one to quantify the predictive power of each dataset as a function of *ρ*. They could be used to inform a decision about which datasets to rely on for predictions in practical settings.

